# Improving sequence-based modeling of protein families using secondary structure quality assessment

**DOI:** 10.1101/2021.01.31.428964

**Authors:** Cyril Malbranke, David Bikard, Simona Cocco, Rémi Monasson

## Abstract

**Motivation:** Modeling of protein family sequence distribution from homologous sequence data recently received considerable attention, in particular for structure and function predictions, as well as for protein design. In particular, Direct Coupling Analysis, a method to infer effective pairwise interactions between residues, was shown to capture important structural constraints and to successfully generate functional protein sequences. Building on this and other graphical models, we introduce a new framework to assess the quality of the secondary structures of the generated sequences with respect to reference structures for the family.

**Results:** We introduce two scoring functions characterizing the likeliness of the secondary structure of a protein sequence to match a reference structure, called Dot Product and Pattern Matching. We test these scores on published experimental protein mutagenesis and design dataset, and show improvement in the detection of non-functional sequences. We also show that use of these scores help rejecting non-functional sequences generated by graphical models (Restricted Boltzmann Machines) learned from homologous sequence alignments.

**Availability:** Supplementary Materials, Data and Code available at https://github.com/CyrilMa/ssqa.

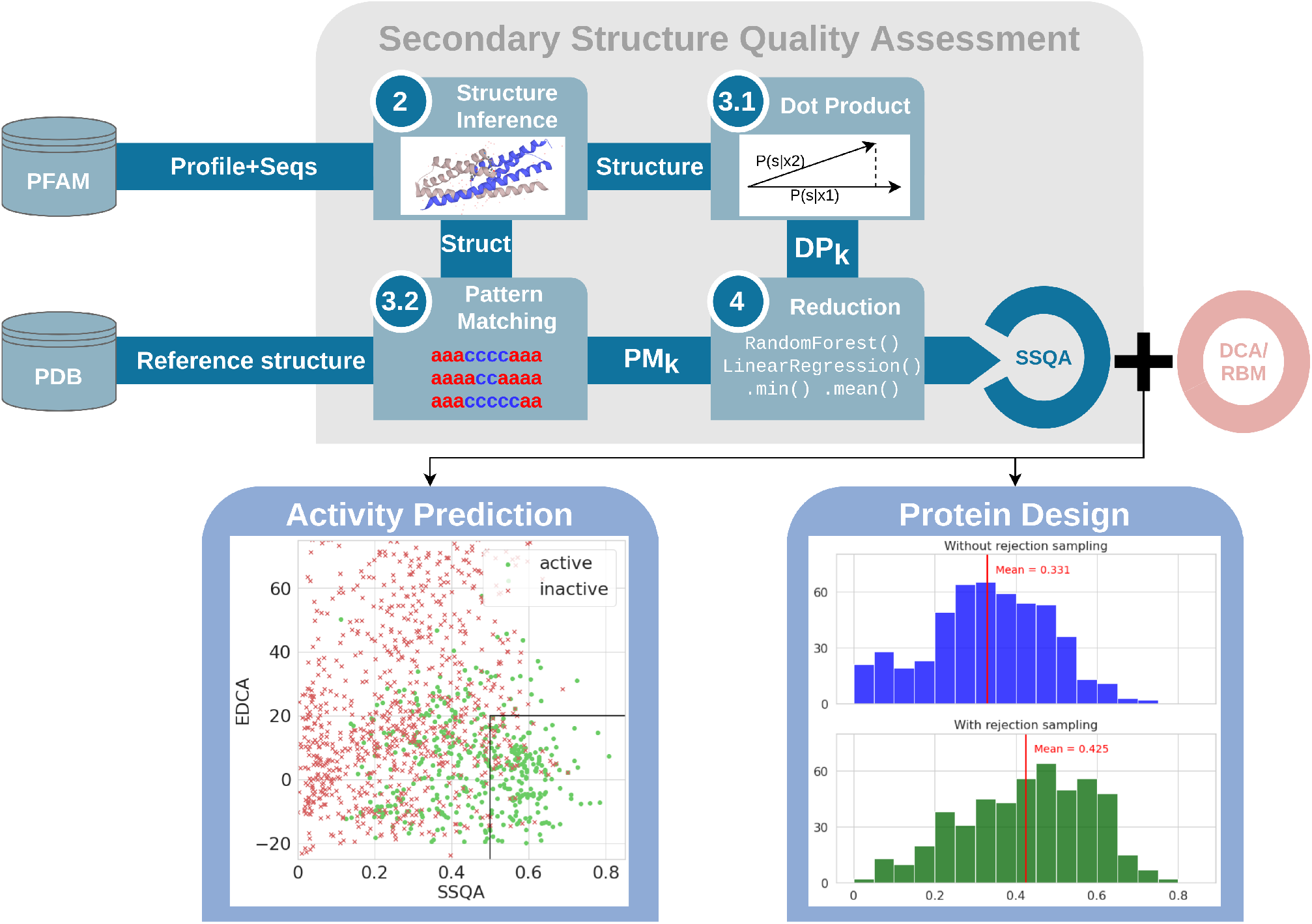

## 1 Introduction

Considerable efforts were devoted over the past decade to the modelling of protein families from homologous sequence data, taking advantage of the tens of millions of available sequences in databases such as Uniprot (The UniProt Consortium (2019)) or PFAM (Bateman *et al.* (2002)). Among sequence-based models, graphical models, in particular Direct Coupling Analysis (DCA), emerged as simple and effective Bayesian inference approaches capturing essential statistical properties of residues in sequence data, such as their conservation and pairwise correlations, see Cocco *et al.* (2018) for a review. DCA outputs a set of statistical pairwise couplings, which are informative about the contact map of the single or multiple folds (Weigt *et al.* (2009), Malinverni *et al.* (2015)) characterizing the family, or about the protein interactions with its partners (Bitbol *et al.* (2016)). In addition, DCA defines a likelihood over the sequence space, which can be used to predict the effects of mutations to a natural sequence in comparison to mutagenesis experiments (Figliuzzi *et al.* (2015), Hopf *et al.* (2017)), or can be sampled to design de novo synthetic proteins, whose viability can be assessed in vivo (Russ *et al.* (2020)).

Despite these successes it remains unclear what aspects of the structural, functional and evolutionary constraints acting on protein sequences are adequately captured by such sequence-based models, and, conversely, what features are inappropriately accounted for. Here, we introduce a method to assess the compatibility of these models with the secondary structure elements common to the family. The goal of our Secondary Structure Quality Assessment (SSQA) method is two-fold. First, we may use SSQA to a posteriori test the validity of sequence-based model predictions, as failure to preserve the secondary structure of a protein is likely to result in a loss of its functionalities. Second, SSQA can be used to guide protein design by helping the production of sequences with adequate secondary structures.

SSQA aims at estimating the similarity between the putative secondary structure associated to a given sequence and a reference structure associated to the protein family. This task is analogous to (tertiary) structure quality assessment, which has received sustained attention in the past years (Derevyanko *et al.* (2018), Baldassarre *et al.* (2020)). Our focus on secondary structure is motivated by several reasons. Secondary structure is known to be largely conserved in protein families (Figure 1), and is therefore a reliable signature of family membership. In addition, state-of-the-art algorithms for secondary structure predictions, such as JPred4 from Drozdetskiy *et al.* (2015), NetSurf0-2.0 from Klausen *et al.* (2018), Wang *et al.* (2016) or Asgari *et al.* (2019) reach very high accuracy levels (85% to 90%). The availability of computationally fast and reliable tools is necessary to the implementation of our quality assessment approach.

**Fig. 1.**
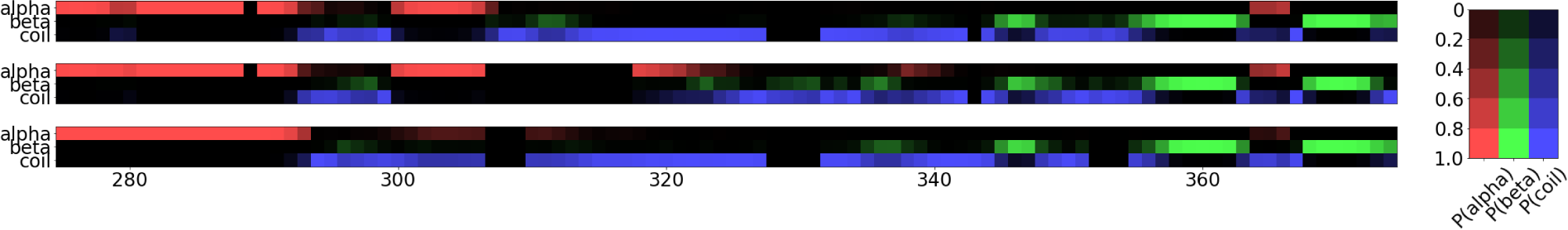
Profiles of predicted secondary structures (*α*-helix, *β*-strand, or coil, see probability values on the right color scale) for three sequences of the betalactamase family (PF00144, aligned sequences : A0A1I2IMA6/54-384, A7SA90/1322-1633, Q18384/50-404). Alignment between the three sequences had a length of 390. For the sake of clarity only positions 275 to 375 are shown. Note the similarities between the three secondary structures.

Our paper is organized as follows. We briefly review graphical models, in particular Restricted Boltzmann Machines, an unsupervised learning framework that encompasses DCA by including high-order couplings between residues in section 2.1, as well as secondary structure inference algorithms in section 2.2. SSQA with its different formulations are presented in section 3. Results on the ability of SSQA to improve functionality/activity prediction are reported in sections 4.1 and 4.2. We then show how protein data-driven design (section 4.3) based on Restricted Boltzmann Machines can be enhanced with SSQA. Conclusive remarks can be found in section 4.3.

## 2 Background

### 2.1 Graphical models for sequence distributions, and Restricted Boltzmann Machines

We will consider hereafter protein sequence distributions ℙ(*x*) expressed by graphical models, where *x* = {*xi*} denotes the sequence of amino acids. A well-known example of graphical model is the so-called Direct Coupling Analysis (DCA), for which

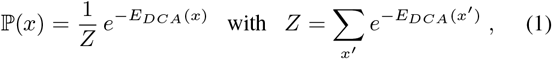

and the energy function is

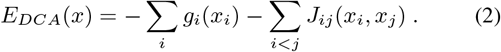

The set of parameters *g_i_*(*x*) and *J_ij_*(*x, y*) are inferred so that the 1- and 2-point statistics, revealing conservation and co-evolution in homologous sequence data match the ones of the model distribution. DCA was shown to be successful for extracting structural information about the 3D conformation of the protein and for designing new functional proteins through the sampling of ℙ(*x*) (see Russ *et al.* (2005) and Russ *et al.* (2020)).

In this work we will consider another class of graphical models called Restricted Boltzman Machine (RBM, see Salakhutdinov (2008) for an overview), which encompass DCA and may also express interactions of order ≥3 between residues in the sequence. RBM were recently shown to be powerful to model amino-acid sequence distributions (Tubiana *et al.* (2019), Bravi *et al.* (2020)). Briefly speaking, RBM are joint probabilistic models on bipartite graphs, with one layer carrying the sequences *x* and another layer the representations *h* = {*h_μ_*}. The energy function for *x, h* is

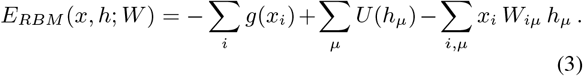

This energy defines the joint distribution of sequences and representations,

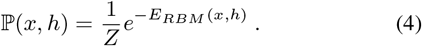

The interactions *W* and the potentials *g*, *U* acting on the input and representation units are trained by maximizing the marginal distribution ℙ(*x*) of the sequences *x* in the dataset. To do this, methods such as Persistent Contrastive Divergence (PCD) can be used (see Tieleman (2008), Tubiana *et al.* (2019)).

Note that the joint probability in (4) also allows one to define the conditional probabilities ℙ(*x|h*) and ℙ(*h|x*). Due to the bipartite nature of the interaction graph these conditional probabilities are factorized, which makes sampling fast and easy. With our choice of a quadratic potential over the representation units, 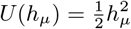, we get the following conditional probabilities for, respectively, representational and sequence units:

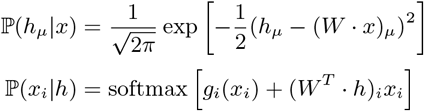

Alternating sampling of the representation and the sequence layers provide an efficient Gibbs procedure to sample P(*x*), see Algorithm 1.

#### Algorithm 1

Gibbs sampling through RBM

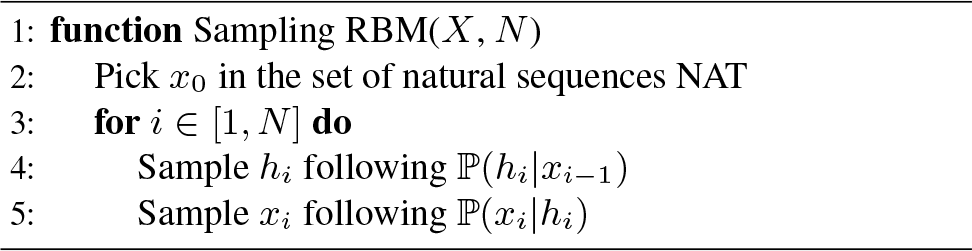

### 2.2 Secondary structure : Definition and Inference

Secondary structure is the three–dimensional form taken by a protein on local scales. The two main secondary structural motifs are *α*-helices (with H-bonds between amino acids that are 3-4 residue apart along the sequence) and *β*-sheets (multiple strands connected by at least 3 H-bonds). We thus represent the secondary structure of a protein by a sequence of a 3-class classification following the primary structure (chain of residues) : *α*-helix, *β*-strands, or “coil” if the residue is part of a disordered segment or an irregular structure. We will also consider more detailed classifications involving 8 classes, see Kabsch and Sander (1983).

Hereafter we consider models, denoted by 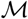, allowing us to estimate the probability for each site *i* in a sequence *x* to be part of a secondary–structure class, e.g. *α*-helix, *β*-strand, or *coil*. These models can be very simple (based on statistics of amino acids), but most successful algorithms now rely on Deep Learning, including 1-dimensional convolutional or recurrent neural networks, in particular LSTM (Hochreiter and Schmidhuber (1997)). Many of these models are proposed in the literature (Asgari *et al.* (2019), Klausen *et al.* (2018)). The most competitive algorithms enrich the sequence of amino acids *x* with hidden Markov model (hmm-er) profiles computed from homologous sequences.

In the present work we focused on: 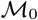, a quick secondary structure predictor based on a 1-dimensional Convolutional Neural Network with 81% accuracy on secondary structure prediction; 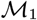, an adapted network based on NetSurfP2-0 from Klausen *et al.* (2018) which reaches 85% accuracy with a relatively light architecture. Both models relied on HMM profiles built through HHsuite (Steinegger *et al.* (2019)).

In Fig. 1, we show the probability maps computed with 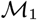 for three sequences of the betalactamase family (PFAM family PF00144). Observation of these profiles on various sequences and families suggests that aligned residues are likely to be part of the same secondary-structure class. In addition, sequences from the same family are likely to have very similar structures, following one or several patterns. Error in predictions are often encountered at the boundary between two distinct classes in the secondary structure; however, even in this case, the probability of the good label generally remains rather high, e.g. 0.4 for the good label and 0.6 for the wrong label.

## 3 Material and Methods

In this section, we propose two ways of assessing the quality of the secondary structure (with respect to a reference secondary structure). Both rely on building a bag of local features from a protein sequence focused on secondary structure. *Dot Product* defines a bag of many raw features (one for each residue of a sequence), quickly computed and fully relying on alignments of the sequences. *Pattern Matching* produces a bag of few refined features (number of secondary structure motifs in the sequence), that require more computation time (quadratic in sequence length) and does not necessarily rely on alignments. Given the refinement of the Pattern Matching features we expect that use of the corresponding features will require less (expensive) annotation data than Dot Product.

### 3.1 Conditional distribution of secondary structures

Let *x* be a protein sequence of length *n*. Its secondary structure *s* is a string of length *n* taking value in 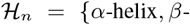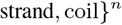, with *s*_*i*_ = *α*-helix*, β*-strand, coil if the residue *i* is part of, respectively, an *α*-helix, a *β*-strand, a disorganized segment (“coil”). In the next part we will also use the DSSP (Kabsch and Sander (1983)) classification with 8 classes of secondary structure motifs.

Let us consider a model 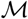 for secondary structure inference from an amino-acid sequence. Given a sequence *x* 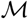 returns *P^x^* ∈ [0, 1]^*n* × 3^ where 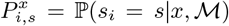 We may then introduce a cost function 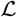 for any secondary structure *s′* in 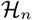,

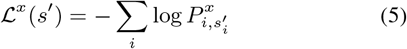

This function is positive and takes low or high values for likely or unlikely secondary structure respectively *x*.

In an approximation in which secondary-structure symbols are independent, this cost function 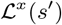 represent the negative log-probability of *x* to have secondary structure 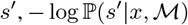. This approximation neglects the presence of correlation between neighbouring sites, but will be use for the sake of simplicity.

### 3.2 Dot Product features

Let us consider for each *i* the probability vector of the secondary structure at residue 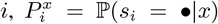. We compare the distribution of probabilities of two sequences *x*_1_ and *x*_2_ at residue *i* through

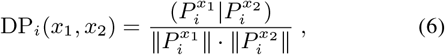

where (·|·) denotes the dot product between two vectors, and ‖·‖ the *L*_2_ norm. DP_*i*_ is a similarity measure between the two secondary structures associated to the sequences *x*_1_ and *x*_2_, equivalent to the cosine of the angle between their two associated vectors. The higher it is the more likely their secondary structures will coincide on site *i*, with 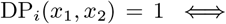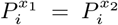. Low values of DP result, on the contrary, from discrepancies between the local predicted secondary structures, *e.g.* DP_*i*_(*x*_1_*, x*2) ≈ 0.105 for 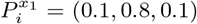 and 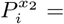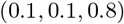

It is also possible to extend the definition of DP above to compare one sequence, say, *x*_1_, to a set of sequences, say, χ (including *N* sequences):

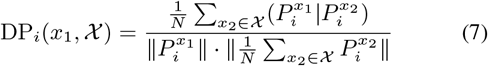

These features will be later referred as the **Dot Product**(DP) features.

### 3.3 Pattern Matching features

A pattern *r* is defined as an ordered set of elements called **motifs**(*r*_*i*_ = (*C_i_, m_i_, M_i_*))_*i<N*_ of ℕ^3^ where *r*_*i*_ is the motif, *C*_*i*_ the motif class (*α*-helix, *β*-strand or coil), *m*_*i*_ and *M*_*i*_ the minimum and maximum size of the motif *r_i_*. *m*_*i*_ and *M*_*i*_ are optional and can be put aside.

A structure *s ∈ (α*-helix*, β*-strand, coil} ^*n*^ is said to **match** the pattern *r* if :

1. ∃(*t_i_*)_*i*≤*N*_ such as *t*_0_ = 0, *t_N_* = *n*
2. ∀*i, m*_*i*_ ≤ *t*_*i*+1_ − *t*_*i*_ ≤ *M*_*i*_
3. ∀*j* such as *t*_*i*_ ≤ *j < t_i_*+1, we have *s*_*j*_ = *C*_*i*_

Afterwards we will denote by *R* the set of secondary structures *s* that match pattern *r*. We define *R*(*x, r*) the probabilistic event of *x* having structure in *R*, and thus matching pattern *r*. We will define Match(*x, r*) the probability of *x* having a structure that matches *r* (realizing event *R*(*x, r*)).

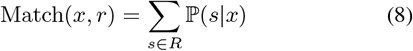

Unfortunately, the computation of the set *R* of structures matching pattern *r* is NP-hard. Brute force computation is not possible, as the size of *R* grows exponentially with the length *n* of the sequence. However, Match(*x, r*) can be calculated in polynomial time, as explained below.

To do so, we make use of the Hidden Markov Model framework. A **Hidden Markov Model**(HMM) is a statistical model in which a system followed or is modeled as a Markov process over a set of hidden (not observable) states, *Z* = (*z_k_*)_*k*_. In addition to this Markov process, there is another process generating observable symbols *Y* = (*y_k_*)_*k*_, where the probability of *y_i_* depends only on *z_i_*.

We consider the following Markov Model :

**Hidden states**: *z_k_* = (*t_k_*_−1_*, t_k_*), the interval of sites [*t_k_*_−1_*, t_k_*[corresponding to motif *r_k_* in class *C_k_*.

**Transition probabilities**: 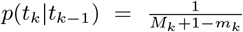 for *t_k_* ∈ [*t_k_*_−1_ + *m_k_, t_k_*_−1_ + *M_k_*], and *p*(*t_k_*|*t_k_*_−1_) = 0 otherwise.

**Observation states**: *y_k_* ∈ 0, 1 where *y_k_* = 1 if a motif matching *r_i_* = (*C_i_, m_i_, M_i_*) conditions is emitted 0 otherwise. We then have *R*(*x, r*) ⇔ ∀*k y_k_* = 1. Given *t_k_*, *t_k_*_−1_, *C_k_* we have *y_k_* emitted following 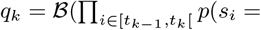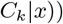 with 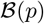 a Bernoulli law of parameter *p*.

The computation of Match(*x, r*) can be done through the sum-product algorithm (Kschischang *et al.* (2001)) which is a dynamic programming method for marginalizing the hidden states in quadratic time similar to the celebrated Viterbi algorithm. More details about the method is available in Supplementary 1. We then obtain Match(*x, r*) and, for each *k*, ℙ(*t_k_*|*R*) and ℙ(*t_k_, t_k_*_+1_|*R*). We define *l_k_* = *t_k_ − t_k_*_−1_ the length of the motif *r_k_* in the secondary structure. These quantities allow us to compute, for each *k*,

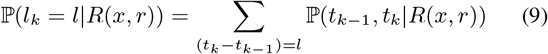

From this we compute our local **Pattern Matching** features, which are the average lengths of secondary structure motifs:

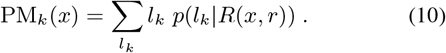

### 3.4 Reduction and full pipeline

We then reduce the bag of features (issued from DP or PM) into a single score able to quantify the quality of the secondary structure. Two approaches are possible, depending on the availability of annotated data.

The first method relies on supervised learning. If experimental measures of the goodness (fitness) of proteins are available, working with Pattern Matching and/or Dot Product features, it is possible to feed these features into a classifier or a regressor for learning with the experimental measures as target. The classifier/regressor used should be adapted to the size of the dataset available but for big enough dataset a Random Forest with 50 trees (from SciKit Learn library, Pedregosa *et al.* (2011)) yielded the best results.

If experimental data are not available we have to rely on “unsupervised” methods. We compute a single score DP from Dot Product features : DP(*x*) = min_*k*_ DP_*k*_(*x, x*_0_) or DP(*x*) = ∑*k* DP_*k*_(*x, x*_0_). And for Pattern Matching : PM(*x*) = min_*k*_ PM_*k*_(*x, x*_0_) or PM(*x*) = ∑*k* PM_*k*_(*x, x*_0_). It is possible to linearly combine both DP and PM into a single score. We experimentally saw that SSQA(*x*) = DP(*x*) + PM(*x*) was already a good approximation of the optimal linear combination of the scores.

After reduction, the pipeline is complete and we obtain a numerical estimate of the quality of the secondary structure of a given sequence. An overview of this pipeline is shown in Figure Fig. 2. Step 2-3 (structure inference, pattern matching and dot product features) have been developed using PyTorch (Paszke *et al.* (2019)), while step 1 relies on HHsuite (Steinegger *et al.* (2019)) and step 4 (reduction) relies on ScikitLearn (Pedregosa *et al.* (2011)) for supervision. PDB structure extraction (Burley *et al.* (2019)) rely on Biotite API (Kunzmann and Hamacher (2018)) in provided repository.

**Fig. 2.**
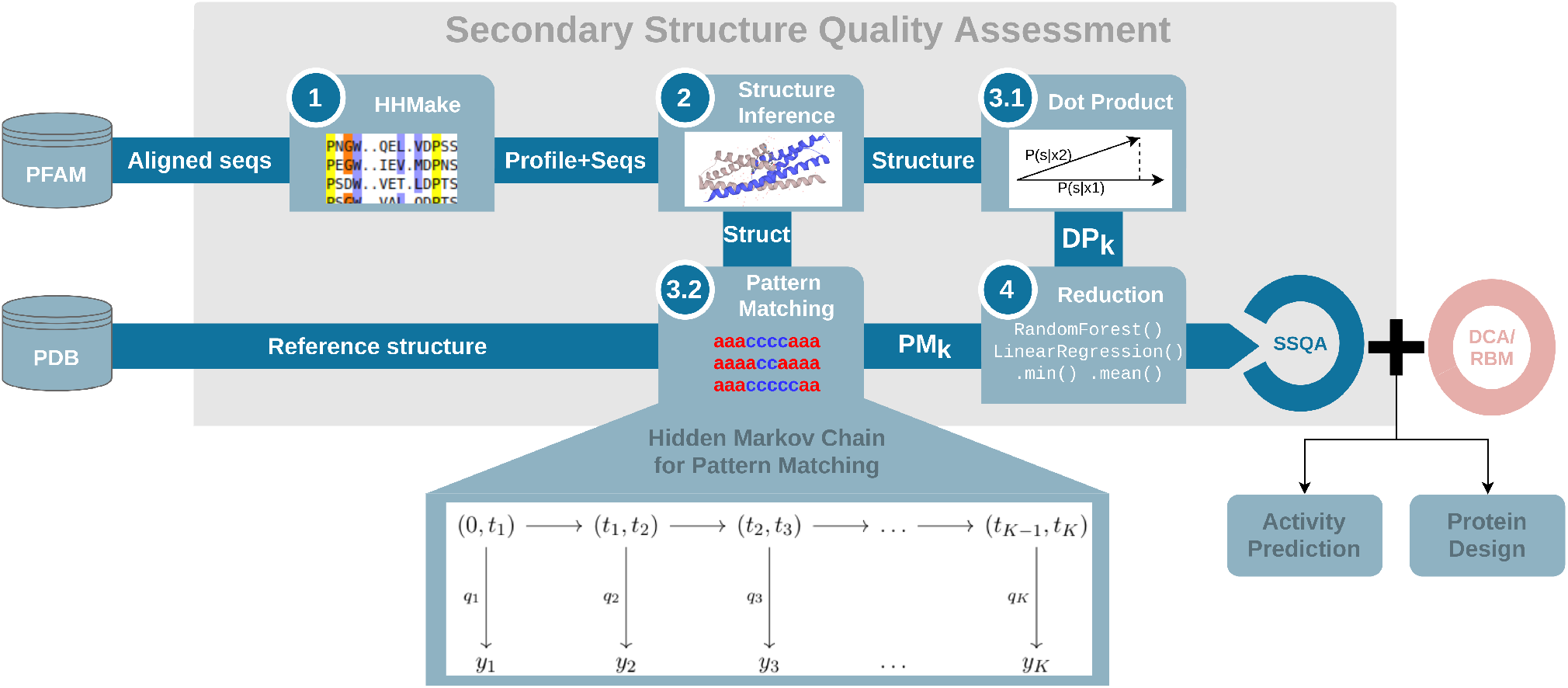
Schema of the secondary structure quality assessment (SSQA) pipeline. First step being the alignment of sequences and computations of the profiles through **HHmake**(image PFAM), then the **structure inference** through a secondary structure predictor (image Uniprot). Then the computation of the **Dot Product** and **Pattern Matching** features with reference structure from PDB and finally the **reduction** through unsupervised or supervised reduction.

## 4 Results

We assess the performance of our approach on existing datasets of various nature. Russ *et al.* (2020) designed new protein sequences from the DCA model learned from homologous sequences of the Chorismate Mutase enzyme (PF07736 in PFAM, alignment referred to as NAT), and measured their fitnesses *in vivo*, see section 4.1. Hopf *et al.* (2017) compiled 23 datasets of mutational effects measured on a collection of proteins (complete list of reference is available in Supplementary 4), see section 4.2.

### 4.1 *A posteriori* screening of DCA-based designed proteins with secondary structure quality assessment

Chorismate Mutase (CM) is an enzyme that catalyzes an intermediate reaction in the synthesis of aromatic amino-acids. Its role in maintaining the balance of these amino acids in the cell is vital, making easy the evaluation of its functionality. In Russ *et al.* (2020), putative protein sequences sampled from DCA model distribution (1), were inserted in E.Coli, in which the CM gene had been removed. The growth rate of these E.Coli presented a bi-modal distribution that allowed for splitting the sequence dataset into *inactive* and *active* samples.

Our objective is to help discriminating active and inactive protein sequences. We will take a look at several scores. First we consider the DCA energy, *E_DCA_*(*x*) in (2) (as available in Russ *et al.* (2020) dataset), which corresponds to the negative log-probability (up to an additive constant) in the Direct Coupling Analysis model. The vast majority of high-energy sequences are inactive, while a substantial fraction of low-energy sequences are active. However, DCA energy alone does not allow for separating active from inactive sequences below some energy threshold, see for instance *E_DCA_ <* 25 in Table 3.c. Russ *et al.* (2020) showed that discrimination performance could be enhanced by a Logistic Regression trained on aligned sequences (MSA) with activity as a target (*MSA Log Reg*). This method could identify generated sequences similar to the one of the test organism (E.Coli) in the low-dimensional space spanned by the top components of the sequence data covariation matrix.

We next study the capability of secondary-structure quality assessment (SSQA) to discriminate between active and inactive sequences, in particular at low *E_DCA_*. To do so we rely on supervised and unsupervised scoring methods based on the Dot Product and the Pattern Matching features. Taking for *x*0 the CM sequence of E.Coli and its secondary structure in PDB (PDB: *1ECM*) as references, we compute the DP and PM features with both 3- and 8-class secondary structures. For the unsupervised scoring functions, we found that 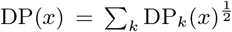 and PM(*x*) = ∑*k* PM_*k*_(*x*) yield the most encouraging results (see Table 3.c). For the supervised scoring functions, we train a model through a Random Forest with 50 trees on the natural sequences (NAT) to target the activity of a sequence and evaluated it on generated sequences (DCA.).

In Fig. 3.a, we plot the DCA energy and the supervised SSQA score (DP+PM) on a same graph with active samples in green and inactive ones in red. We see that SSQA helps discriminate active and inactive samples with low energy. Most of the sequences with low energy have been correctly labelled as inactive by SSQA, while most samples with good SSQA and low DCA energy are active. In the low *E_DCA_*/high SSQA domain delimited by the black lines in the figure 85% of the sequences are active.

**3.a.**
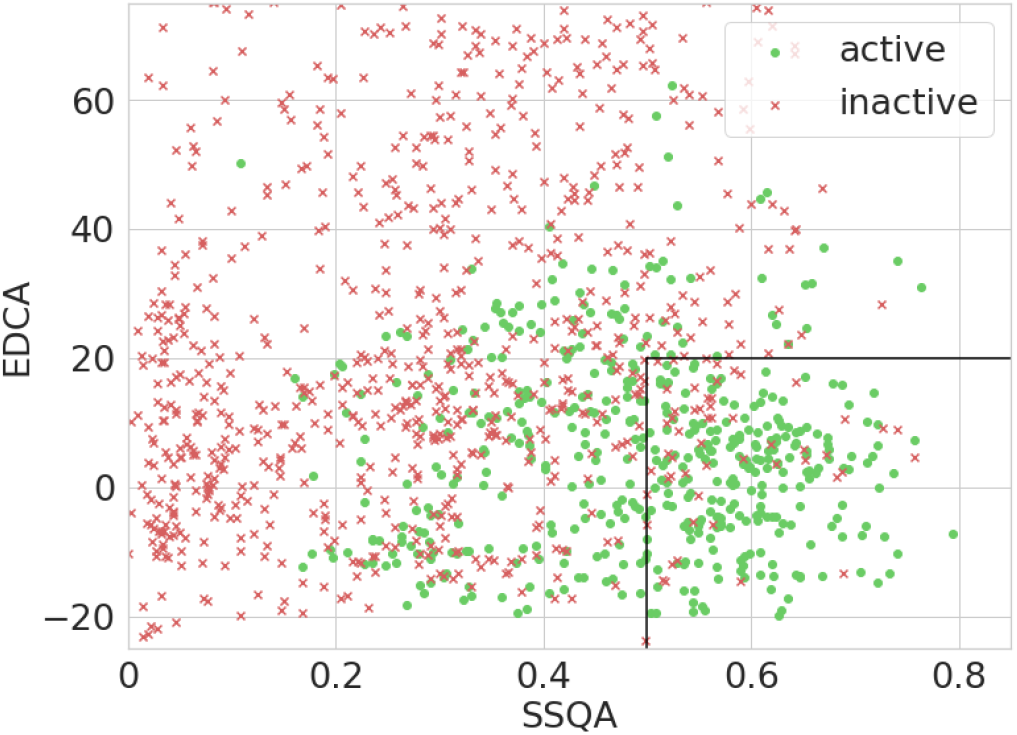
*E_DCA_* and SSQA of generated samples. Green dots are the active samples. As we can see, the combination of DCA energy and SSQA allow a good discrimination. 82% of the samples in the black block are active.

**3.b.**
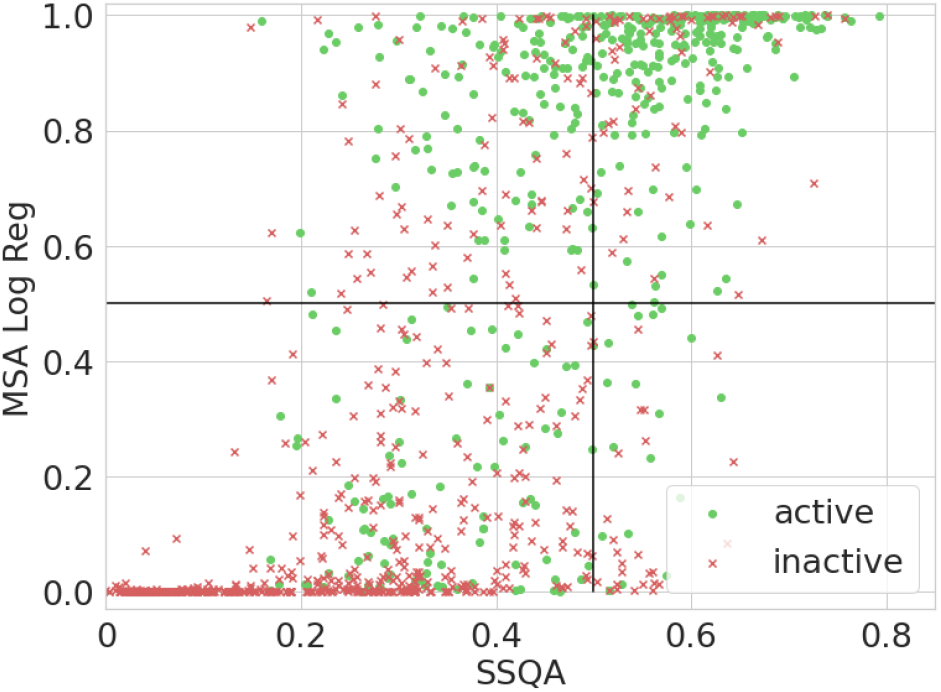
MSA Log Reg from Russ *et al.* (2020) and SSQA of generated samples. Green dots are the active samples. MSA Log Reg is performing Logistic Regression of the MSA of the sequences with activity as a target. The Spearman correlation between the two scores is *ρ* = 0.65.

**3.c.**
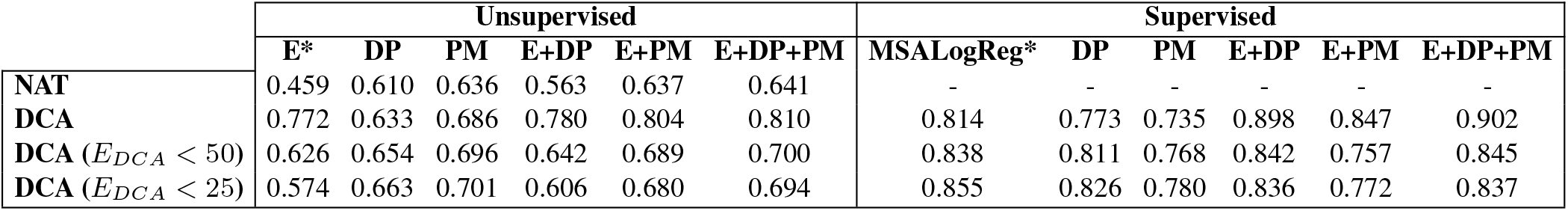
AUROC for inactive samples detection for different combination of features, *MSA Log Reg* and *E_DCA_* from Russ *et al.* (2020) dataset, Pattern Matching (PM) features and Dot Product features (DP) with and without supervision. We computed the AUC with the activity as a target for different datasets : natural sequences (NAT), generated sequences (DCA) and subset of the generated sequences by focusing on low energy samples (*E_DCA_ <* 25) or on samples generated with different sampling temperatures *T* (the higher *T*, the more the samples will be generated with freedom towards the training set). As we can see the use of SSQA features (DP, PM) has a particular interest on low energy samples that are indistinguishable from natural sequence from statistics of order 1 and 2. The combination DCA energy and SSQA features often yield a very good discrimination of low activity samples (AUC : 0.914 with supervision and 0.810 without on the full dataset). ROC curves associated to displayed AUCs are available in Supplementary 2, with plotted correlation between displayed scores and experimental activity.

In Fig. 3.b, we show the scatter plot of *MSA Log Reg* and of the supervised SSQA score (DP+PM). The high value of the Spearman coefficient underlines the correlations between the enrichment method developed by Russ *et al.* (2020) and secondary structure features. This correlation may either reflect a causal effect, *i.e.* preservation of secondary structure is a key ingredient to the functionality of the protein, or simply that the similarities at the secondary structure level are indicative of the phylogenetic similarities in the CM family.

Last of all we notice in Table 3.c that DP clearly outperforms PM when used with supervision, while PM is better without supervision. This is an important remark to take into consideration, since, for the many studies that do not rely on experimental measurements of protein viability, supervised methods to improve the precision of SSQA cannot be used.

### 4.2 Secondary structure quality assessment on mutational datasets

We generalize the method on mutational datasets extracted from multiple mutagenesis studies compiled in Hopf *et al.* (2017). Each of these datasets contain sequences with generally one or few mutations around a wild-type sequence, with the experimentally determined values of their in vitro or in vivo fitnesses. Hopf *et al.* (2017) perform mutational effect predictions through DCA couplings, see (2), that we take as baseline for our own predictor.

To quantify the performance of secondary structure quality assessment, we train models with features computed through DP and PM through cross validation. The secondary-structure patterns are retrieved from PDB when available, or inferred with the Pattern Matching inference method (see Supplementary 1). The model we select is a Random Forest Regressor (50 trees) fitted with the experimental fitnesses as targets through cross validation. We then linearly combine Dot Product and Pattern Matching scores with the DCA score (energy of mutated sequence) from Hopf *et al.* (2017), weighting of the scores are optimized through Linear Regression and cross-validation. We compute for each obtained score the Spearman correlation *ρ* with the ground-truth experimental measurements. A selection of these correlations can be found in Fig. 4.a. Fig. 4.b shows the scatter plot of the Spearman correlations obtained with the DCA coupling estimates (E) only vs. the ones where DCA couplings are combined with Dot Product and Pattern Matching (E+DP+PM). We see that, for most datasets, both Dot Product and Pattern Matching bring an improvement to the mutational effect prediction.

**4.a.**
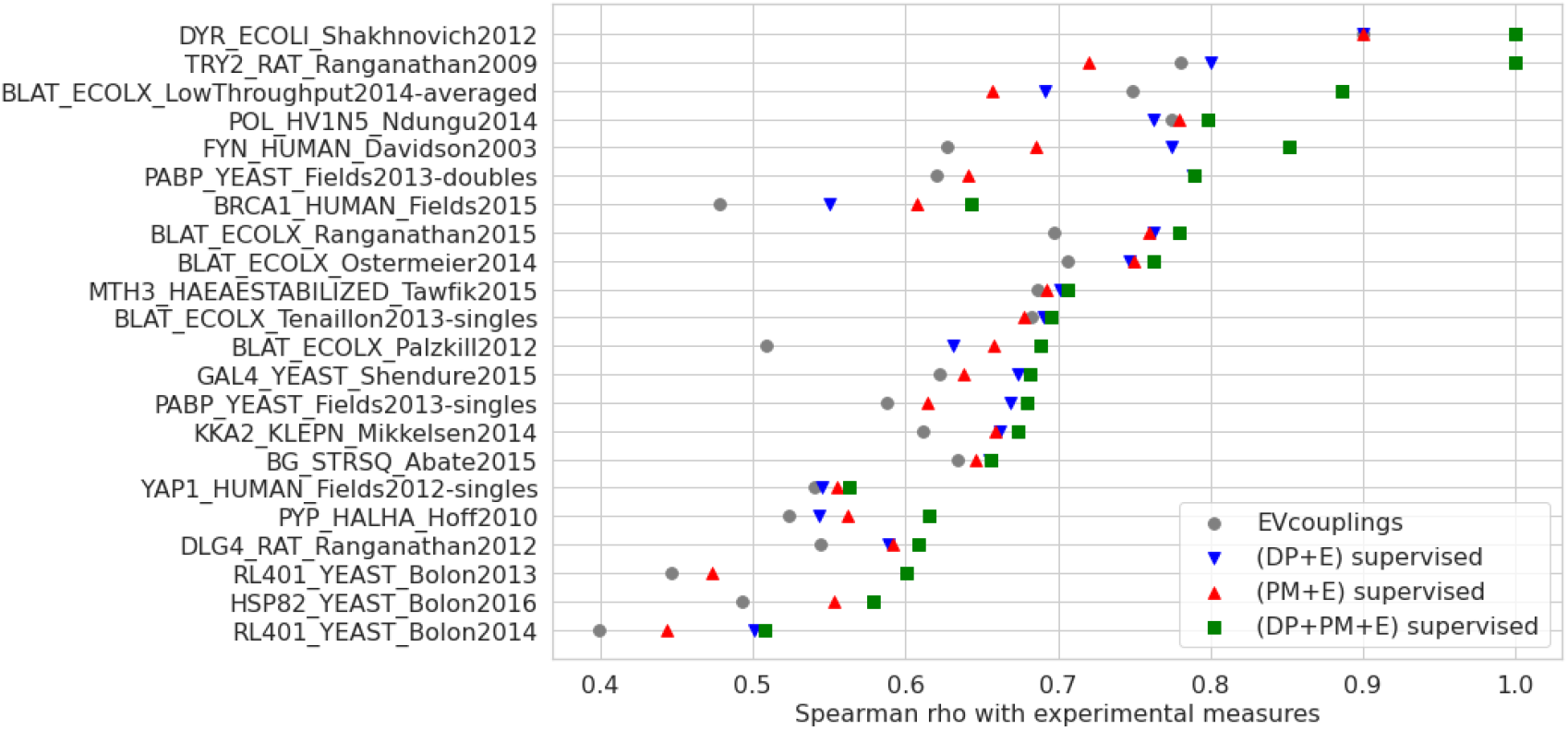
Spearman correlations between experimental fitness measurements computed through cross validation with DCA energy (E) from Hopf *et al.* (2017), Dot Product (DP) and Pattern Matching (PM) features, and their combinations, for different datasets.

**4.b.**
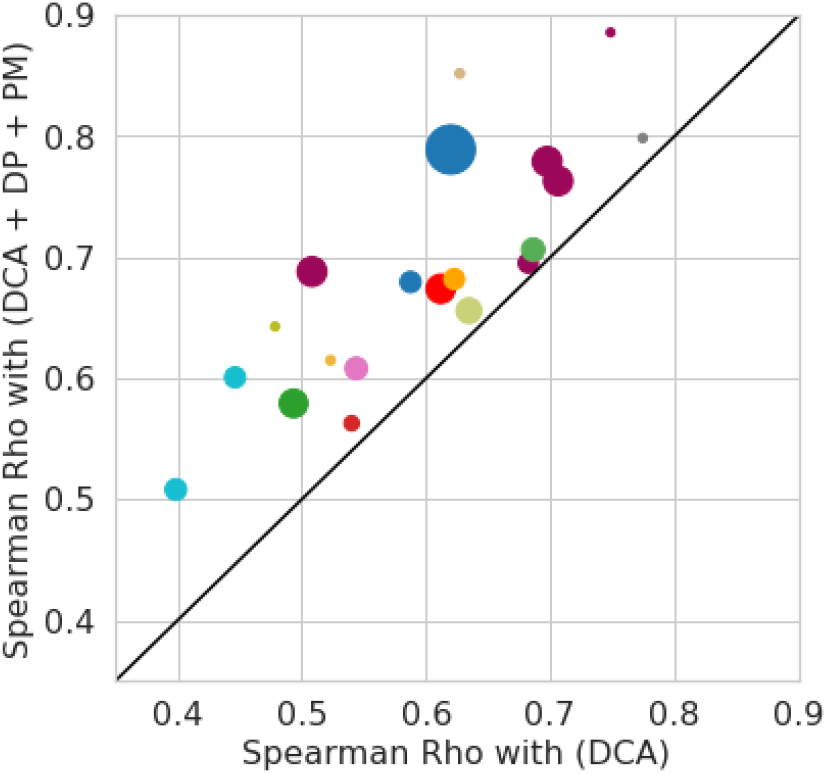
Scatter plot of the Spearman correlations *ρ* between the experimental measurements of fitness and the scores combining DCA energy, as well as DP and PM features (E+DP+PM) vs. DCA energy only (E) over the different datasets. The sizes of the dots are proportional to the sizes of the corresponding datasets. The improvement brought by SSQA is non-negligible in particular for bigger datasets.

**4.c.**
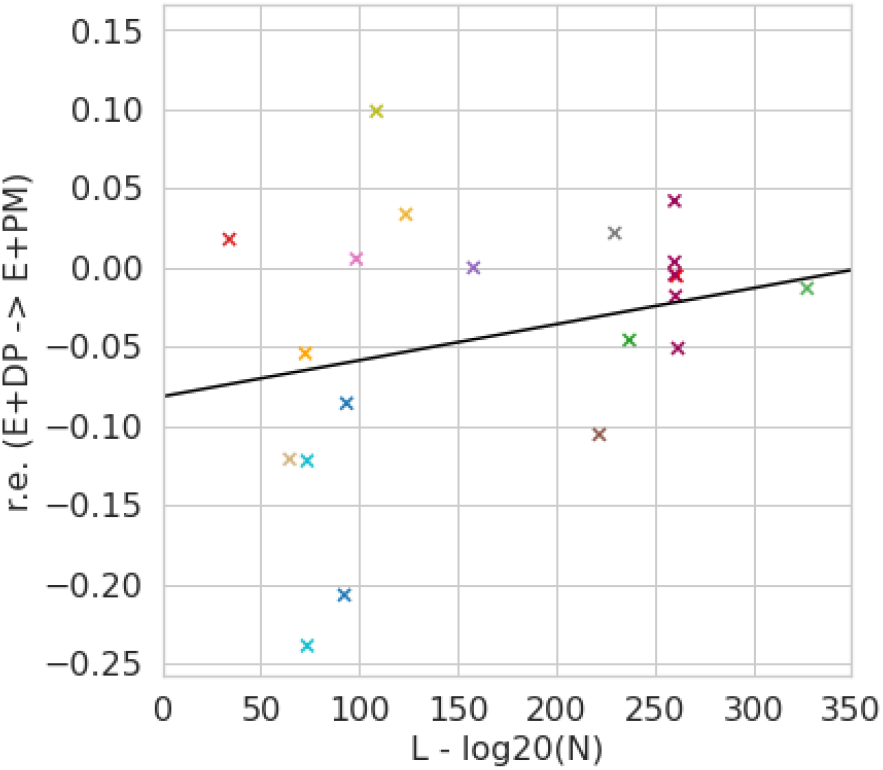
Relative enrichment from Dot Product method to Pattern Matching given the relative size of the dataset *N* compared to the size the sequence *L−* log 20(*N*). As the quantity of available annotated data decreases Dot Product become less and less performing compared to Pattern Matching.

In Fig. 4.c, we show the relative improvement 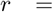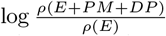 as a function of the ratio *L* − log_20_(*N*), where *L* is the length of the reference sequence and *N* the size of the dataset. We observe a slight correlation between *r* and *L* – log_20_(*N*) (Pearson *ρ* = 0.32, p value = 0.14). Dot Product often outperforms Pattern Matching for large training dataset, whereas Pattern Matching is superior to Dot Product for small datasets. The few sophisticated features produced by Pattern Matching are helpful for long proteins or small datasets, but it is advisable to use the many raw features of Dot Product when sequences are short and numerous.

### 4.3 Improved Restricted Boltzmann Machine-based sequence sampling with secondary structure sequence assignment

We now ask whether SSQA can improve graphical-model-based generating models. We consider a RBM energy *ERBM* (*x*) for sequence *x*, and a scoring function *m*(*x, r*) of the secondary-structure quality (local or global) of sequence *x* with respect to pattern *r*. We propose to sample sequences following the distribution

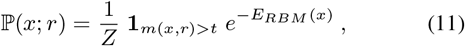

where *Z* is a normalization constant, and the indicator function **1** rejects all sequences with scores lower than the threshold *t*.

In practice, we sample the RBM distribution of (4) with Gibbs sampling, see Algorithm 2. Then we perform rejection sampling as following to simulate ℙ(*x*; *r*).

#### Algorithm 2

Gibbs rejection sampling through RBM

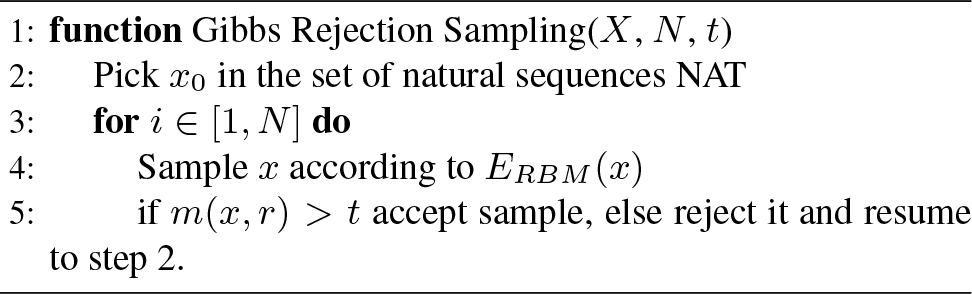

Using Chorismate Mutases dataset, we train a Restricted Boltzman Machine from the NAT alignment of PFAM (PF07736) and Persistent Contrastive Divergence. The “*L*_1_*b*” normalization defined in Tubiana *et al.* (2019) is used. The hidden layer is composed of 200 Gaussian units. After training, the RBMs are used to sample 2000 sequences by Gibbs sampling (30 steps). Rejection with different thresholds are performed, to enforce harder and harder secondary structure requirement. We also sample from an independent-site model (based on 1-point statistics over residues only) as a baseline.

We take the unsupervised pattern matching score to reject samples, as this score requires knowledge about the secondary structure only, thereby making the method applicable to proteins for which no experimental data is available. We use the supervised combined E+DP+PM score to assess the quality of our generated samples. Even though methods used for rejection and for functionality assessment are similar, the latter still gives a good idea of the improvement brought by the rejection. We find that the fraction of sequences predicted active in the generated dataset are 26% and 51% with, respectively, no (*t* = − ∞) and high (*t* = 0.65) rejection. Results are displayed in Fig. 5, where a clear shift toward good structures samples is observed as threshold *t* is increased.

**Fig. 5.**
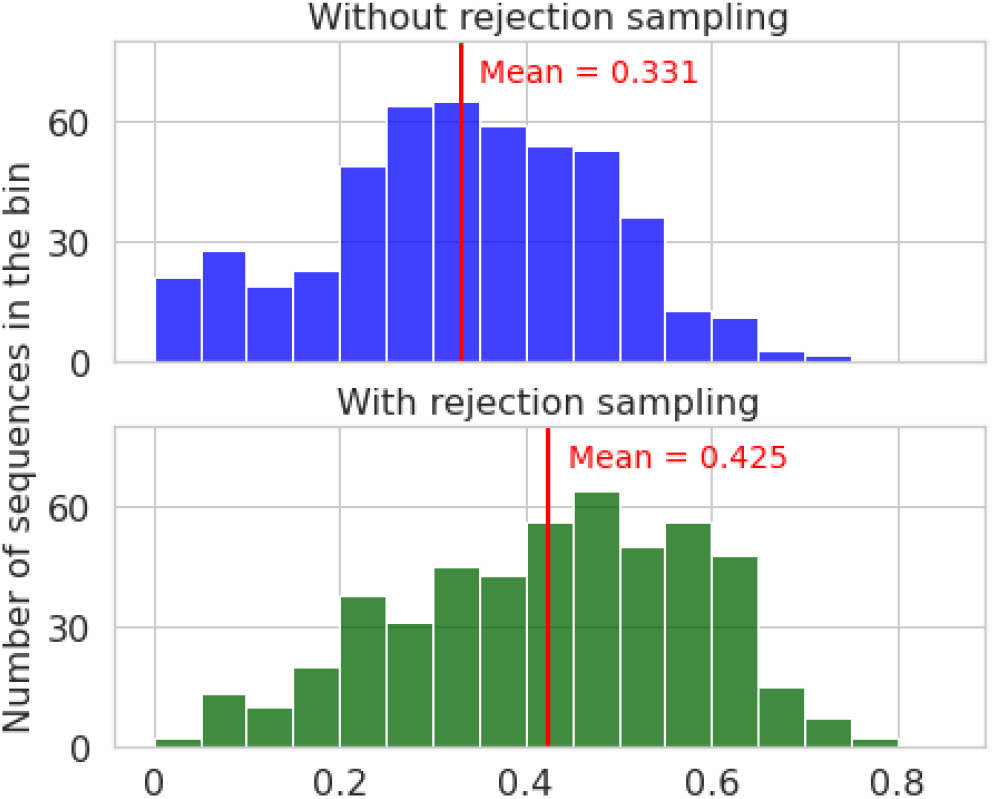
Activity likelihood distribution, assess with supervised E+DP+PM predictor, of sequences generated by RBM without and with high rejection sampling (with unsupervised SSQA), out of 500 sequences generated by the RBM. The numbers of putatively active sequences is much higher with hard rejection (green) than with no (blue).

## Conclusion

In summary, we have proposed multiple scoring functions for assessing the compatibility of protein sequences with respect to a reference secondary structure. Our approach is computationally tractable, has intuitive meaning, and shows promising performance. *A posteriori* validation of sequences generated in previous works shows the ability of our method to detect dysfunctional proteins, and constitutes an improvement compared to standard graphical model-based methods. These results strongly suggest that quality assessment is a practical way to exploit secondary structure, despite the ~15% error rate of the best available secondary structure prediction algorithms.

The results we report on Chorismate Mutases or on mutational effect datasets showed a great complementarity between secondary structure quality assessment and direct coupling analysis; for instance, functionality of sequences with both good SSQA and good DCA energy have been shown to be very high (more than 80% for Chorismate Mutases). It is not surprising that DCA or RBM captures functionally relevant information on residues (for instance, at or close to binding sites) beyond secondary structure alone. However, it is less clear what statistical features of the sequence data are overlooked by graphical models, and, yet, essential to secondary structure prediction. A possible answer to this question can be found in Supplementary 3. We show, for the betalactamase family (see Majiduddin *et al.* (2002)), that SSQA brings particular improvement in the activity prediction task for mutations happening on *β*-strand, where DCA models are failing to yield good prediction. While it may not be inconceivable that accounting for some *β*-motifs may require a complex pattern of couplings, beyond what DCA can accommodate for, further systematic studies are required to understand the origin of the complementarity between their scores.

In addition, we observed that SSQA methods, in particular Pattern Matching, may lead to enhanced performance with little sequence annotation *e.g.* additional structural information, or little experimental data. Indeed, state-of-the-art secondary structure prediction software have been tested and validated on huge datasets, including all protein families with known structures. Use of these methods for SSQA of sequences attached to a single protein family may therefore be seen as an illustration of knowledge transfer. In this context, addition of tertiary structure quality assessment method such as in Baldassarre *et al.* (2020) would be interesting for further developments.

Last of all, the efficient computation of SSQA scores reported in this work suggest other applications and their integration at the heart of sequence generative processes, such as sampling with rejection, as done here. Furthermore, it would be interesting to integrate these scores into reward functions for Reinforcement Learning process or loss functions for Neural Networks, in generative networks (such as Repecka *et al.* (2019), Hawkins-Hooker *et al.* (2020)) or representation network (Rives *et al.* (2019), Alley *et al.* (2019)), which is made possible by their differentiability.

## Acknowledgements & Fundings

SC and RM were supported by the Agence Nationale de la Recherche grant numbers ANR-17-CE30-0021 RBMPro and ANR-19-CE30-0021 Decrypted. CM is recipient of a PhD funding from AMX program, Ecole Polytechnique and benefits from financial support from the Centre de Recherche Interdisciplinaire (CRI) through “Ecole Doctorale Frontières de l’Innovation en Recherche et Education – Programme Bettencourt”. The authors declare no conflict of interest.

