## Supplementary Materials for "Improving sequence-based modeling of protein families using secondary structure quality assessment"

### Supplementary Material for *Improving sequence-based modeling of protein families using secondary structure quality assessment*

Cyril Malbranke, David Bikard, Simona Cocco and Rémi Monasson

January 30, 2021

#### Contents

**1 Pattern Matching details** **2**
1.1 Likelihood of a pattern . . . . . . . . . . 2
1.2 Pattern Inference . . . . . . . . . . 3

**2 Supplementary figures about *A posteriori screening of DCA-based designed proteins with SSQA*** **5**

**3 Improvements of SSQA in function of the secondary structure in the betalactamase** **7**

**4 References to Datasets used for "Secondary structure quality assessment on mutational datasets"** **9**

### 1 Pattern Matching details

A pattern $r$ is defined as an ordered set of elements called **motifs** $(r_i = (C_i, m_i, M_i))_{i<N}$ of $\mathbb{N}^3$ where $r_i$ is the motif, $C_i$ the motif class ($\alpha$-helix, $\beta$-strand or coil), $m_i$ and $M_i$ the minimum and maximum size of the motif $r_i$. $m_i$ and $M_i$ are optional and can be put aside.

A structure $s \in \{\alpha\text{-helix}, \beta\text{-strand}, \text{coil}\}^n$ is said to **match** the pattern $r$ if :

1. $\exists (t_i)_{i \leq N}$ such as $t_0 = 0$, $t_N = n$
2. $\forall i,\ m_i \leq t_{i+1} - t_i \leq M_i$
3. $\forall j$ such as $t_i \leq j < t_{i+1}$, we have $x_j = C_i$

Afterwards we will denote by $R = \{s \in \mathcal{P}(\{\alpha\text{-helix}, \beta\text{-strand}, \text{coil}\}^n)\}$ the set of secondary structures that match the pattern $r$. We will define $\text{Match}\, M(x, r)$ the probability of $x$ having a structure that matches $r$

$$\text{Match}(x, r) = \sum_{s \in R} \mathbb{P}(s|x) \tag{1}$$

Unfortunately, the computation of $R$ the set of structures matching pattern $r$ is NP-hard. We will not be able to compute this set that grows exponentially with the size $n$ of the sequence. However, we found a way to compute $\text{Match}\, M(x, r)$ in polynomial time.

#### 1.1 Likelihood of a pattern

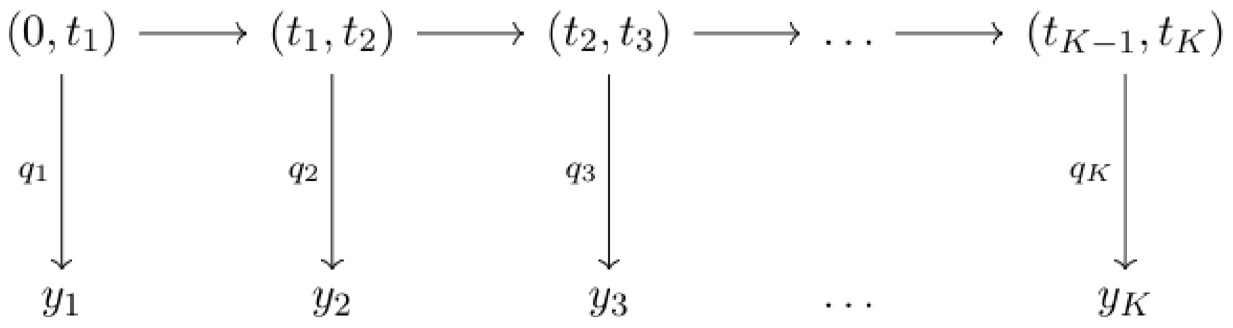


Figure 1: Hidden Markov Model with $z_i = (t_{i-1}, t_i)$ for Pattern Matching prediction

Given the probability matrix of the structure $P^x$ defined previously we want to assess the probability of this probabilistic structure to match $r$, denoted by $\mathbb{P}(R(x, r))$.

It is possible to represent this problem with a Hidden Markov Model. A **Hidden Markov Model** (HMM) is a statistical model in which a system followed or is modeled as a Markov process $Z = (z_k)_k$, where $Z$ is not observable. In addition of this Markov process, there is another process $Y = (y_k)_k$, observable and such that $\forall i$, $y_i$ depends only on $z_i$. The objectives we can meet with these model are multiple : decoding (finding the most likely $z$), marginalizing (finding $p(z_i|y_1, ..., y_k)$) ...

In our case we consider the following Markov Model :

**Hidden states** : $z_k = (t_{k-1}, t_k)$ the intervals of residues $[t_{k-1}, t_k[$ of the motif $r_k$ of class $C_k$.

**Transition probability** : $p(t_k|t_{k-1}) = \frac{1}{M_k - m_k}$ for $t_k \in [t_{k-1} + m_k, t_{k-1} + M_k]$ and $p(t_k|t_{k-1}) = 0$ otherwise.

**Observation states** : $y_k \in \{0,1\}$ where $y_k = 1$ if a motif matching $r_i = (C_i, m_i, M_i)$ conditions was emitted. given $t_k$, $t_{k-1}$, $C_k$ it is easy to see that $y_k \sim q_k = \mathcal{B}(\prod_{i\in[t_{k-1},t_k[} p(s_i = C_k))$

We can see from the previous definition that $R(x,r) \iff \forall k, y_k = 1$. We then have :

$$\text{Match}(x,r) = \mathbb{P}(R(x,r)) = p(y_1 = 1, \dots, y_K = 1)$$

In order too compute this probability, we will rely on our Markov chain. We here recall a dynamic programming way of marginalizing a Hidden Markov Model with a process called **sum product algorithm** (see [13]). For a HMM model with observations $y_k$ and hidden states $z_k$ we define recursively :

$$\alpha_{k+1}(z_{k+1}) = p(y_{k+1}|z_{k+1}) \sum_{z_k} p(z_{k+1}|z_k)\alpha_k(z_k)$$
$$\beta_k(z_k) = \sum_{z_{k+1}} p(y_{k+1}|z_{k+1})p(z_{k+1}|z_k)\beta_{k+1}(z_{k+1})$$

we then have after computation :

$$\alpha_k(z_k)\beta_k(z_k) = p(z_k, y_0, \dots, y_K)$$
$$\sum_{z_k} \alpha_k(z_k)\beta_t(z_k) = p(y_0, \dots, y_K)$$

We will be using the sum-product algorithm with our own Hidden Markov Model (Figure 1). After re-arrangement, it gives us :

$$\alpha_{k+1}(t_{k+1}) = \sum_{t_k} p(s_{[t_k,t_{k+1}[} = C_k)p(t_{k+1}|t_k)\alpha_k(t_k)$$
$$\beta_k(t_k) = \sum_{t_{k+1}} p(s_{[t_k,t_{k+1}[} = C_k)p(t_{k+1}|t_k)\beta_{k+1}(z_{k+1})$$

In this case:

$$\begin{aligned}\text{Match}(x,r) = \forall k,\ \mathbb{P}(R(x,r)) &= p(y_1 = 1, \dots, y_N = 1)\\ &= \sum_{t_k} \alpha_k(t_k)\beta_k(t_k)\end{aligned}$$

And in particular :

$$\begin{aligned}\text{Match}(x,r) = \mathbb{P}(R(x,r)) &= p(y_1 = 1, \dots, y_K = 1)\\ &= \alpha_K(n)\beta_K(n) = \alpha_K(n)\end{aligned}$$

#### 1.2 Pattern Inference

For a lot of proteins (in particular the ones that are driving attention), structures from which we infer patterns are available online, we retrieve most of them on the Protein Data Bank [3]. For others we may be require to infer the pattern from the result of our secondary structure prediction.

We will expand our Hidden Markov Model so the class of each motif is integrate in the hidden state. We will have to also adapt our transition probability to our new model (see Figure 2). The adapted model will then be :

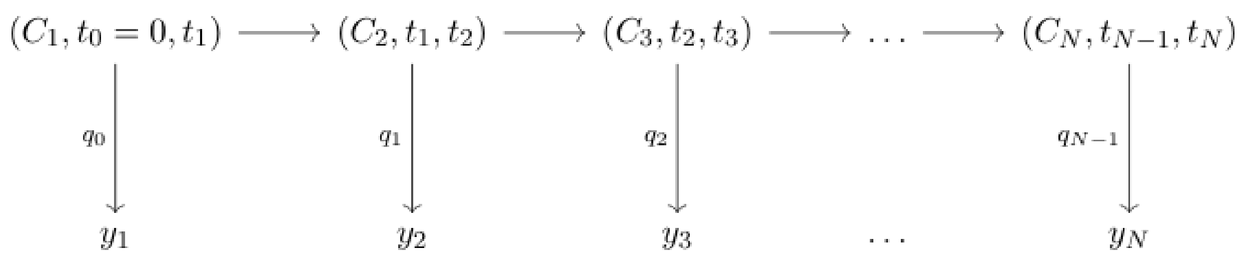


Figure 2: Hidden Markov Model with $z_i = (C_i, t_{i-1}, t_i)$ for Pattern Matching Inference

**Hidden states** : $z_k = (C_k, t_{k-1}, t_k)$ the class $C_k$ of the motif $r_k$ the intervals of residues $[t_{k-1}, t_k[$ of the motif $r_k$ of class $C_k$.

**Transition probability** : The transition probability will be model like
$p(C_k, t_{k-1}, t_k | C_{k-1}, t_{k-1}, t_{k-2}) = p(C_k | C_{k-1}) \cdot p(l_k = t_k - t_{k-1} | C_k, t_{k-1})$

Where $p(C_k|C_{k-1})$ the probability of having a motif of class $C_k$ following a motif of class $C_{k-1}$ and $p(l_k|C_k)$ the probability of a motif $C_k$ having a length $l_k$ are inferred from available training dataset.

From a technical point of view since we don't know a priori the length of the motif it is necessary to add a final stationary state $S$ with $p(C_k = S|C_{k-1} = S) = 1$ and $p(l_k = 0|C_k = S) = 1$.

With this formalism we can run the max-product algorithm (or Viterbi algorithm) to find the most likely pattern given the predicted secondary structure.

#### 2 Supplementary figures about *A posteriori screening of DCA-based designed proteins with SSQA*

Here on Figure 3, we plot the ROC curve for both unsupervised and supervised metrics. As we can see unsupervised scores, even though of course they perform less than supervised scores are able to perform some discrimination of active and inactive samples thus potentially helping improving sampling

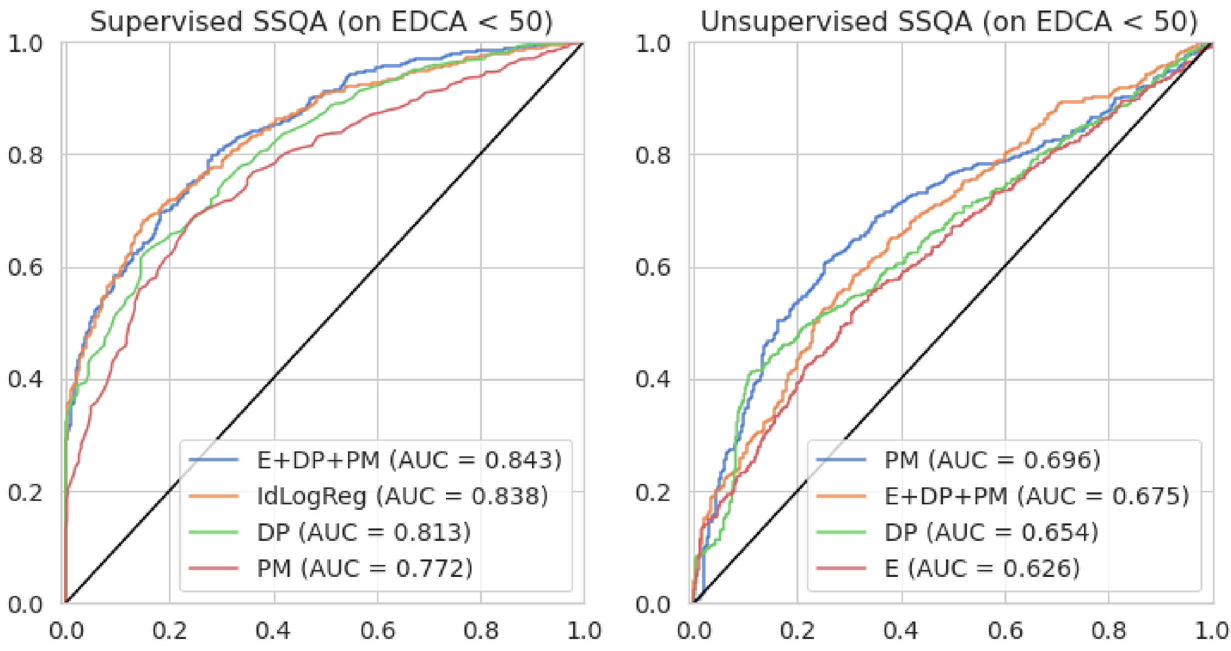


Figure 3: ROC Curve for inactive samples detection with unsupervised and supervised SSQA scores. We focused on low energy samples statistically equivalent to natural samples in terms of statistics of order 1 and 2

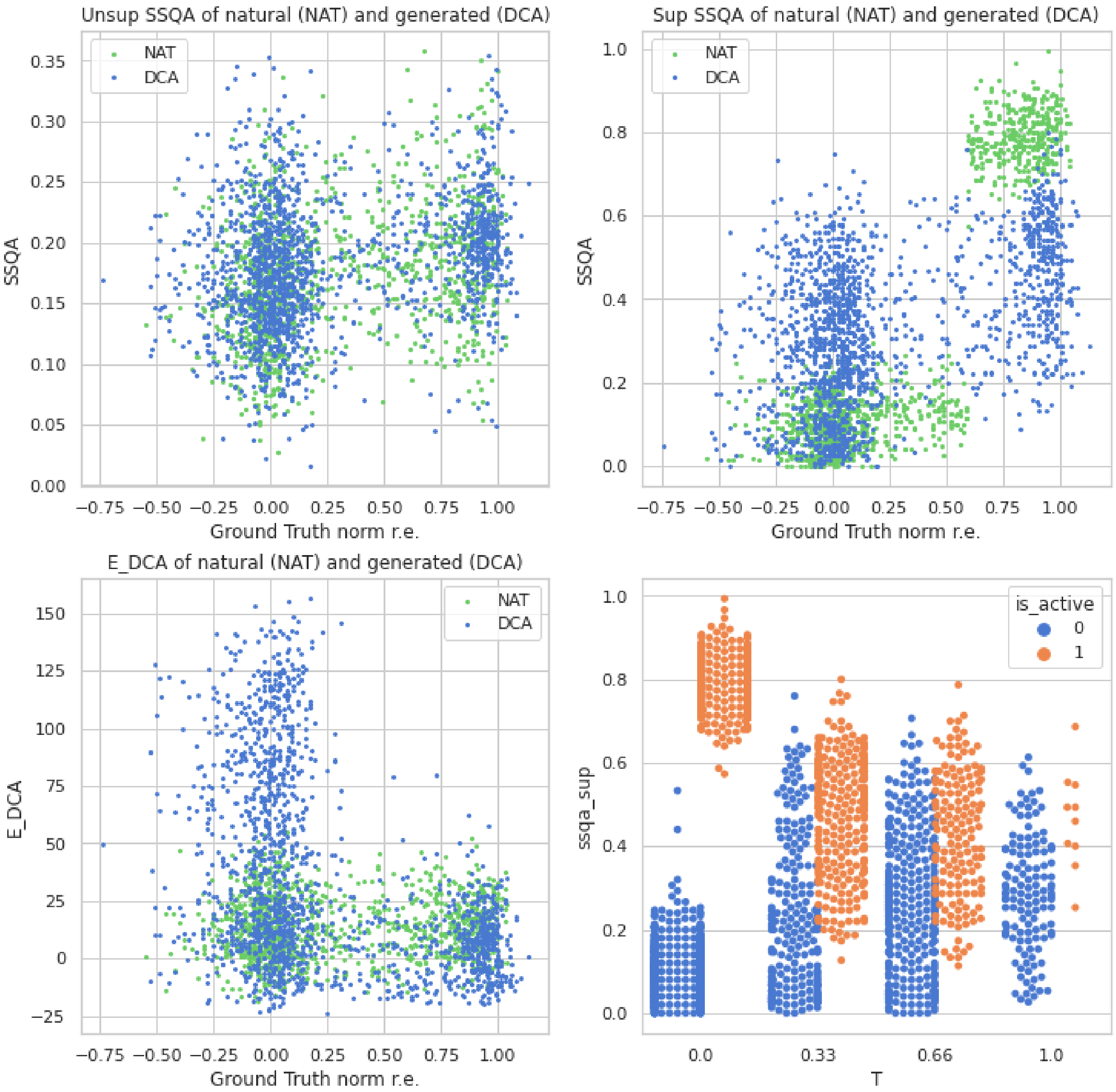


Figure 4: Plot of *unsupervised* SSQA, *supervised* SSQA and $E_{DCA}$ of generated and natural sequences in function of the exprimental activity (Around 0 being inactive, around 1 being active). As we can see, DCA is able to discriminate easily a lot of bad sequences but fails for some of them. *Supervised* SSQA is also very able to discriminate some bad sequences as well as well *unsupervised* SSQA though it is less visible. Last figure is the violinplot in function of the temperature of generation. 0 being Natural Sequence and the higher T the more liberty will be taken with natural sequences. *supervised* SSQA shows a good discrimination at every level.

### 3 Improvements of SSQA in function of the secondary structure in the betalactamase

We work with single mutations dataset sequence from beta lactamase [4] (Uniprot ID : P62593). Activity for each single mutation have been experimentally determined (see Figure 5). We linearly combined Dot Product and Pattern Matching features we computed with taking DCA energy from [10], and built activity predictors out of these metrics. As we can see Figure 6 or Figure 7 with for 8-class, predictor based on DCA energy was able to reach 70% to 75% accuracy on $\alpha$-helix and coil but was failing on $\beta$-strand. SSQA predictor brought important improvements in particular on $\beta$-strand

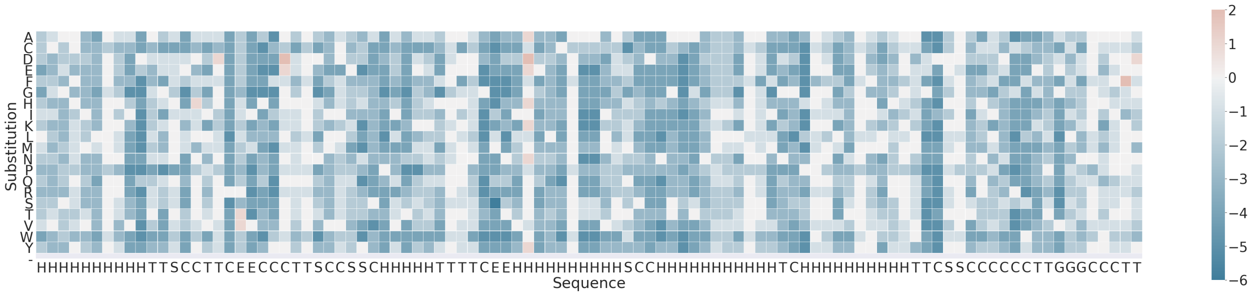


Figure 5: Experimental Mutation Effects on a segment of the beta-lactamase (structure in x-axis), red squares show increasing in activity, blue dots show decreasing activity

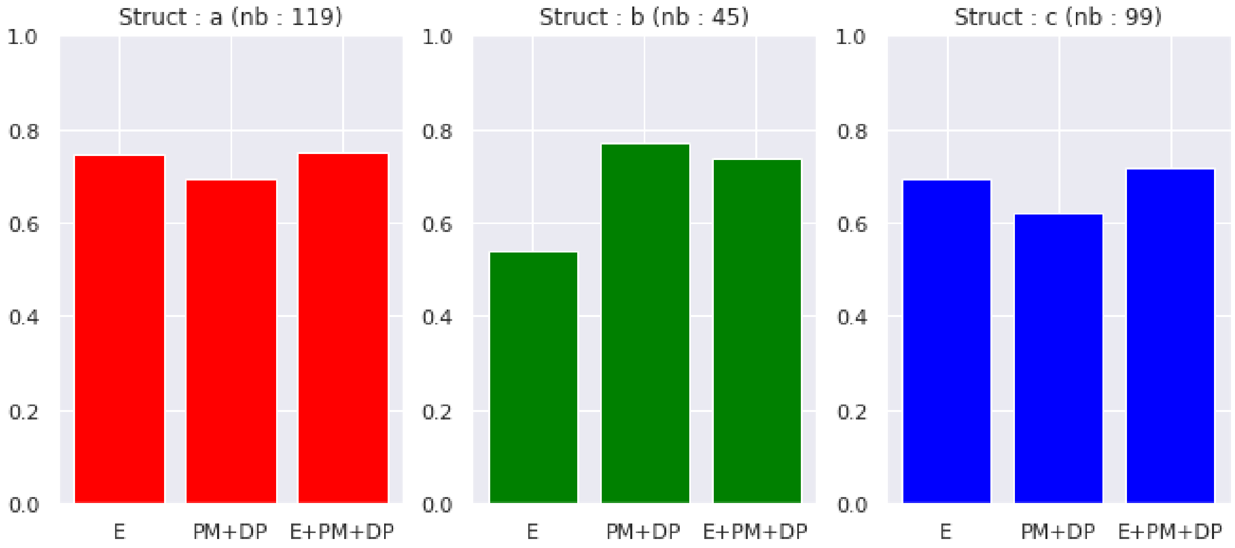


Figure 6: Balanced accuracy for activity prediction on Beta-lactamase for single mutations given the 3-class secondary structure of the mutated residue. As we can see SSQA brings particular improvement on $\beta$-strands where interaction between residues are usually more complex than in $\alpha$-helix.

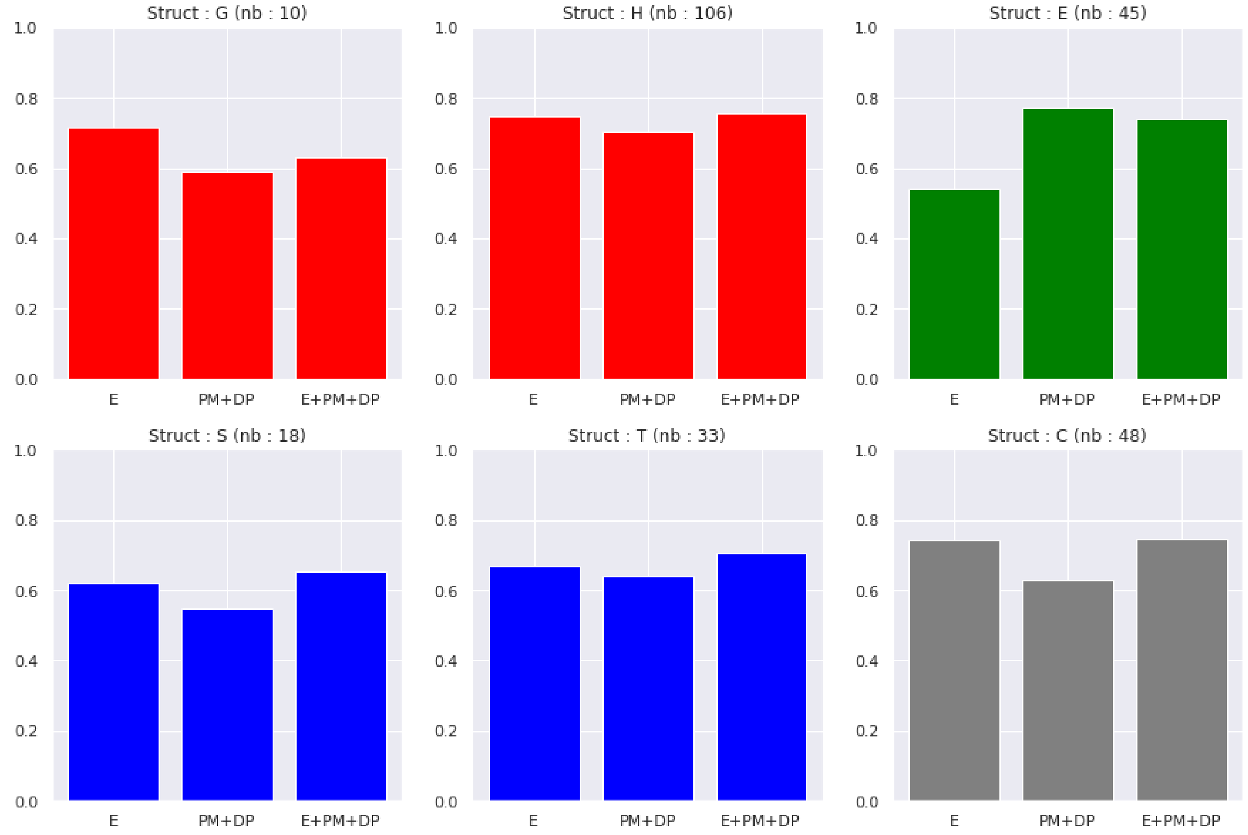


Figure 7: Balanced accuracy for activity prediction on Beta-lactamase for single mutations given the 8-class secondary structure (only 6 classes are present in the structure) of the mutated residue. As we can see SSQA brings particuluar improvement on $\beta$-strands where interaction between residues are usually more complex than in $\alpha$-helix.

### 4 References to Datasets used for ”Secondary structure quality assessment on mutational datasets”

Here is a list of dataset collected in Hopf et al. [10] and use in section 4. :

| ID | Reference |
|---|---|
| POLG_HCVJF_Sun2014 | Qi et al., PLOS Pathogens 2014 [20] |
| UBE4B_MOUSE_Klevit2013-singles | Starita et al., PNAS 2013 [25] |
| PA_FLU_Sun2015 | Wu et al., PLOS Genetics [28] |
| RL401_YEAST_Bolon2014 | Roscoe et al., JMB 2014 [23] |
| PABP_YEAST_Fields2013-singles | Melamed et al., RNA 2013 [16] |
| GAL4_YEAST_Shendure2015 | Kitzmann et al., Nat Methods 2015 [12] |
| RL401_YEAST_Bolon2013 | Roscoe et al., JMB 2013 [24] |
| PABP_YEAST_Fields2013-doubles | Melamed et al., RNA 2013 [16] |
| HG_FLU_Bloom2016 | Doud & Bloom, Viruses 2016 [7] |
| DLG4_RAT_Ranganathan2012 | McLaughlin et al., Nature 2012 [15] |
| BG_STRSQ_Abate2015 | Romero et al., PNAS 2015 [22] |
| BLAT_ECOLX_Palzkill2012 | Deng et al., JMB 2012 [5] |
| BLAT_ECOLX_Ostermeier2014 | Firnberg et al., Mol Biol Evol 2014 [8] |
| HSP82_YEAST_Bolon2016 | Mishra et al., Cell Reports 2016 (in press) [18] |
| BLAT_ECOLX_Ranganathan2015 | Stiffler et al., Cell 2015 (Table S1 and S4) [27] |
| BRCA1_HUMAN_Fields2015 | Starita et al., Genetics 2015 (Table S2) [26] |
| KKA2_KLEPN_Mikkelsen2014 | Melnikov et al., NAR 2014 [17] |
| YAP1_HUMAN_Fields2012-singles | Araya et al., PNAS 2012 [1] |
| MTH3_HAEAESTABILIZED_Tawfik2015 | Rockah-Shmuel et al., PLOS Comp Bio 2015 (File S3) [21] |
| PYP_HALHA_Hoff2010 | Philip et al., PNAS 2010 (Table S1) [19] |
| BLAT_ECOLX_LowThroughput2014-averaged | Firnberg et al., Mol Biol Evol 2014 [8] |
| FYN_HUMAN_Davidson2003 | Di Nardo et al., JMB 2003 (Table 2 and 3) [6] |
| DYR_ECOLI_Shakhnovich2012 | Bershtein et al., PNAS 2012 (Tables 1, S1, S2) [2] |
| POL_HV1N5_Ndungu2014 | Mann et al., PLOS Comp Bio 2014 (Supplementary Table 1, 2) [14] |
| TRY2_RAT_Ranganathan2009 | Halabi et al., Cell 2009 (Table S2) [9] |
| BLAT_ECOLX_Tenaillon2013-singles | Jacquier et al., PNAS 2013 (Supplementary Data 1) [11] |

[10] Thomas Hopf, John Ingraham, Frank Poelwijk, Charlotta Schärfe, Michael Springer, Chris Sander, and Debora Marks. Mutation effects predicted from sequence co-variation. *Nature Biotechnology*, 35, January 2017.

[25] Lea M. Starita, Jonathan N. Pruneda, Russell S. Lo, Douglas M. Fowler, Helen J. Kim, Joseph B. Hiatt, Jay Shendure, Peter S. Brzovic, Stanley Fields, and Rachel E. Klevit. Activity-enhancing mutations in an E3 ubiquitin

ligase identified by high-throughput mutagenesis. *PNAS*, 110(14):E1263–E1272, April 2013. Publisher: National Academy of Sciences Section: PNAS Plus.

[26] Lea M. Starita, David L. Young, Muhtadi Islam, Jacob O. Kitzman, Justin Gullingsrud, Ronald J. Hause, Douglas M. Fowler, Jeffrey D. Parvin, Jay Shendure, and Stanley Fields. Massively Parallel Functional Analysis of BRCA1 RING Domain Variants. *Genetics*, 200(2):413–422, June 2015. Publisher: Genetics Section: Communications.

[27] Michael A. Stiffler, Doeke R. Hekstra, and Rama Ranganathan. Evolvability as a function of purifying selection in TEM-1 -lactamase. *Cell*, 160(5):882–892, February 2015.

[28] Nicholas C. Wu, C. Anders Olson, Yushen Du, Shuai Le, Kevin Tran, Roland Remenyi, Danyang Gong, Laith Q. Al-Mawsawi, Hangfei Qi, Ting-Ting Wu, and Ren Sun. Functional Constraint Profiling of a Viral Protein Reveals Discordance of Evolutionary Conservation and Functionality. *PLOS Genetics*, 11(7):e1005310, July 2015. Publisher: Public Library of Science.
